# Functional RNA Interference in *Hymenoscyphus fraxineus:* Hairpin RNA-Induced Gene Silencing Of A Polyketide Synthase-like Gene

**DOI:** 10.64898/2026.02.02.703346

**Authors:** Linus Hohenwarter, Arvid Hanke, Alexandra Bassler, Gitta Langer, Gabi Krczal, Veli Vural Uslu

**Affiliations:** Translational Epigenetics, RLP Agroscience GmbH, Breitenweg 71, D-67435 Neustadt-Mussbach, Germany; Department of Forest Protection, Northwest German Forest Research Institute (NW-FVA), Grätzelstraße 2, D-37079 Göttingen, Germany; Centre for Organismal Studies (COS) Heidelberg, University of Heidelberg, INF230, D-69117 Heidelberg, Germany

**Keywords:** *Hymenoscyphus fraxineus*, ash dieback, RNA interference, small RNA, gene silencing, translational inhibition

## Abstract

European ash dieback caused by the invasive ascomycete species *Hymenoscyphus fraxineus* poses the most prominent danger to common ash trees (*Fraxinus excelsior*) in Europe. The disease is widely distributed in Europe and currently no efficient management strategy is available. Host-induced gene silencing and exogenous dsRNA applications have shown great potential for controlling fungal diseases in crop plants. In this study, we reported *in silico* evidence for the presence of a functional RNA interference pathway in *Hymenoscyphus fraxineus*. Moreover, we showed that the transgenic expression of a double stranded RNA (dsRNA) leads to inhibition of translation of its target polyketide synthase-like gene, a fungal endogene. We explored whether the dsRNA could be introduced exogenously and demonstrated that *H. fraxineus* can take up externally applied dsRNA molecules. This study highlights the RNA interference mechanism in *H. fraxineus* and suggests exoRNA applications as a promising approach to control European ash dieback.

## 1. Introduction

Ash dieback, caused by the ascomycete fungus *Hymenoscyphus fraxineus*, has led to an extreme decline in common ash (*Fraxinus excelsior*) in Europe with a loss of up to 85% (Coker et al. 2019). *H. fraxineus* infects ash leaves by airborne ascospores, colonising the petiole and growing further back inside the xylem, leading to necrosis of surrounding tissue and the characteristic *dieback* of branches (Gross et al. 2014). After abscission, the *Chalara* anamorph emerges on infected petioles, releasing spermatia that fertilize an ascogonium to form new apothecia in summer releasing infectious ascospores (Gross et al. 2012). Additionally, the disease is associated with stem collar necrosis, where the primary infection with *H. fraxineus* weakened ash trees for secondary fungal infections leading to increased mortality and high risk of downfall (Peters et al. 2023). The fungus, native to east Asia, was first detected in symptomatic ash trees in north-western Poland in the early 1990s (Przybył 2002; Kowalski 2006) but recent findings from older herbarium samples suggest an introduction at least 14 years earlier (Agan et al. 2023). The fungus is now widely spread across Europe (Enderle et al. 2019), reaching northern Spain and southern Italy (Stroheker et al. 2021; Migliorini et al. 2022), effectively overlapping with the entire native habitat of ash (Beck et al. 2016). To date, there are no effective plant protection measures to stop an infection or provide preventive protection.

RNA interference (RNAi) is an essential pathway for the development of new plant protection strategies. RNAi is a multilayered regulatory mechanism present in almost all eukaryotic organisms with common functions in regulating endogenous genes, securing genome stability by silencing transposons and repetitive elements, and defence against invading nucleic acids (Lax et al. 2020). In fungi, RNAi is also involved in pathogenesis (Gaffar et al. 2019). The mechanistic core of RNAi is conserved among all organisms, but the regulatory details diverged significantly on a large evolutionary scale. RNAi is initiated by double stranded RNA (dsRNA) derived from endogenous (e.g., MIR genes, tandem repeats, antisense transcripts) or foreign (e.g. virus replication, transgene) sources (Torres-Martínez and Ruiz-Vázquez 2017). dsRNA is cleaved into small interfering RNA (siRNA) by DICER-like endonucleases (DCL) that range between 21-25-nt, depending on the organism and DCL variant involved. siRNAs are incorporated into ARGONAUTE proteins (AGO), and various proteins are recruited to form regulatory complexes, such as the RNA-induced silencing complex (RISC) or RNA-induced transcriptional silencing (RITS) complex. A growing number of different siRNA classes have been described in fungi and other organisms, facilitating the diversity of regulatory mechanisms (Lax et al. 2020). Gene silencing is achieved post-transcriptionally (PTGS) by translationally blocking or degrading the target transcript (RISC), or transcriptionally (TGS) by inducing DNA methylation and/or heterochromatin formation at the target locus (RITS) (Chang et al. 2012). In both pathways, target recognition is mediated by sequence homology of the AGO-incorporated siRNA. In plants, the silencing signal gets amplified by RNA-dependant RNA polymerase (RDR) that synthesizes dsRNA from certain single-stranded RNA (e.g. aberrant transcripts or cleavage products) resulting in a positive-feedback loop and the formation of secondary siRNA. Homologues of plant RDRs are also found in fungi, but their role in RNAi varies between fungal species. The function of RDRs in the basal fungus *Mucor circinelloides* is comparable to that in plants in amplifying the silencing signal, but secondary siRNA formation has not been documented in filamentous fungi such as *Neurospora crassa, Fusarium asiaticum*, or *Verticillium dahlia* (Torres-Martínez and Ruiz-Vázquez 2017; Mann et al. 2023). In *N. crassa* and other filamentous fungi, RDRs are involved in response to DNA damage and tandem repeat silencing, where dsRNA is generated from aberrant transcripts, and additionally, RDRs mediate genome stability by silencing unpaired DNA in meiosis.

This study aims to investigate the mechanisms of RNAi-driven gene silencing in *H. fraxineus*, which is a promising approach for controlling ash dieback.

## 2. Materials and methods

### 2.1 Fungal strains and culture conditions

*Hymenoscyphus fraxineus* strain NW-FVA 1856 was isolated from a naturally infected *Fraxinus excelsior* stem collar necrosis in northern Schleswig-Holstein, Germany (Waldgehege Fahrenstedthof, mark 24860, Böklund, Abt. 3410a) in 2013 (Langer 2017). Fungal cultures were grown in liquid or solid MYP-medium (7 g malt extract, 0.5 g yeast extract, 1 g peptone from soybean, and 15 g micro-agar for solid medium in 1 l ddH2O, pH 5.8) at constant 22 °C without direct illumination. In the same growing chamber, fluorescent LEDs at a 16/8 h light/dark cycle illuminated other shelves.

### 2.2 Vector construction pSilent HfPKS-IR

PKS-like genes in *H. fraxineus* were identified using *Ascochyta rabiei* PKS1 (ACS74449.1) (Akamatsu et al. 2010) protein sequence as the query in a tBLASTn search (https://blast.ncbi.nlm.nih.gov/Blast.cgi) against *H. fraxineus* genome assembly (BioProject Accession: PRJEB14441, Assembly: GCA_911649665.1). One PKS-like gene (CAG8950773.1, HYFRA_00002986) shared sequence similarity (Coverage: 96%, Identity: 55%, E-value: 0.0) and was selected as a target for gene silencing.

The hairpin RNA construct comprised of the partial 287 bp sequence complementary to the *Hymenoscyphus fraxineus* PKS-like gene. A 117 bp spacer sequence from the *Solanum lycopersicum* RDR intron 3 (Solyc05g007510) separated the sense and antisense orientation of the PKS fragment. The addition of *Sph*I-*Eco*RI (consecutive on 5’ end) and *Hind*III restriction sites flanking the PKS fragment completed the 709 bp PKS-IR construct. The complete fragment was synthesized by Invitrogen’s GeneArt Gene Synthesis service. The DNA fragment was delivered as a plasmid (pMK_RQ_HfPKS-IR). Ligation of a *Sph*I/*Hind*III released PKS-IR fragment into *Sph*I/*Hind*III opened pSilent-1 (Nakayashiki et al. 2005) resulted in plasmid pSilent HfPKS-IR (Supplementary Fig. S1).

### 2.3 Transformation of fungal protoplasts

The generation of protoplast from *Hymenoscyphus fraxineus* mycelium and PEG-mediated DNA transfer was done as described earlier (Lutz et al. 2023) with some modifications. Briefly, one 7 mm mycelium plug inoculated 50 ml liquid MYP and grown statically for 7 days prior to rejuvenation by mixing twice for 10 sec in a blender (Blender 8011EB, Waring, Torrington, Connecticut, USA) and diluting 25 ml blended culture with an equal volume of fresh MYP medium. After 3 days, approximately 600 mg mycelium was harvested by centrifugation (10 min, 2800 x *g*) and washed twice with 50 ml sterile ddH2O. The washed pellet was mixed with 10 ml enzyme solution (1 M MgSO_4_, 50 mM tri-sodium citrate, 1,75% w/v Driselase™ Basidiomycetes sp., 1,75% w/v lysing enzymes, pH 5.8) and incubated 20 h at 30 °C with gentle shaking at 50 rpm. Both enzymes were purchased from Sigma-Aldrich (St. Louis, Missouri, USA). Using a 100 µm sieve, protoplasts were separated from undigested mycelium and adjusted to 15 ml with MgSO4 buffer (1 M MgSO4, 50 mM tri-sodium citrate, pH 5.8). The suspension was mixed carefully with 15 ml 850 mM sucrose and overlayed with 1200 µl STC buffer (500 mM sucrose, 10 mM Tris-HCl, 50 mM CaCl2, pH 8.0). After centrifugation (15 min, 2800 x *g*), an equal volume of floating protoplasts was recovered and 600 µl protoplast suspension was gently mixed with linearized plasmid (10 – 20 µg in 30 µl) and incubated for 20 min at RT. 1.5 ml PEG solution (40% w/v PEG4000 in STC) was added, mixed gently, and incubated for another 20 min at RT. Then, 5 ml of MYP_Reg_ (500 mM sucrose in MYP, pH 5.8) was added and regenerated in the dark for 3 days. Regenerated protoplasts were mixed with 45 ml solid MYPReg (45 °C), split into two petri dishes (Ø 90 mm), and incubated for 4 days in the dark. The cultures were overlayed with 25 ml solid MYP containing 200 µg/ml hygromycin B. Emerging transformants were transferred to solid MYP plates containing 100 µg/ml hygromycin B.

### 2.4 Screening for transgenic lines

Genomic DNA was extracted from mycelium of a 7-day-old liquid culture as described in (Cenis 1992). Stable integration of the DNA was verified by standard PCR using DreamTaq Green DNA Polymerase (Thermo Fischer Scientific). Amplification of the full-length PKS-IR insert was not possible, probably due to the formation of secondary structures in the amplicon. Therefore, strands were amplified separately with primers for sense (pSilent1_insert_fw and Spacer_rv) and antisense (pSilent1_insert_rv and Spacer_fw). The presence of a hygromycin resistance cassette was verified using Hygro_fw and Hygro_rv. Primer sequences are listed in (Supplementary Table S1).

### 2.5 RNA extraction and expression analysis

Fungal mycelium was grown for 7 days on a small (0,7 – 1 cm^2^) filter paper that was homogenized in liquid nitrogen. Total RNA was extracted using the QIAzol lysis reagent (Qiagen, Venlo, Netherlands) and subsequently, DNaseI treated (New England Biolabs, Ipswich, Massachusetts, USA) for 15 min at 37 °C and purified using the RNeasy MinElute Cleanup Kit (Qiagen). cDNA was synthesized from 0.25 – 1 µg RNA with oligo-dT primers included in the ProtoScript First Strand cDNA Synthesis Kit (New England Biolabs). Expression levels of PKS and reference gene UBC (CAG8960749.1) were analysed in technical triplicates by quantitative Real-Time PCR (qRT-PCR) using the Luna Universal qPCR Master Mix (New England Biolabs) and CFX96 Real-Time System with a C1000 Touch thermal cycler and Bio-Rad CFX Maestro 1.1 Software Version 4.1.2433.1219 (Biorad, Hercules, California, USA). Relative expression changes were calculated using the 2^-ΔΔct^ method. Primer sequences are listed in (Supplementary Table S1).

### 2.6 RISC extraction and small RNA sequencing

RISC complexes (AGO proteins associated with siRNA) were extracted from ground mycelium (as above) with the TrapR Kit (Lexogen GmbH, Vienna, Austria) described in Grentzinger et al. 2020. Eluted small RNAs were provided to Lexogen for library preparation and sequencing on Illumina NextSeq P2 at 100 cycles. Adapter-trimmed and demultiplexed reads were analysed with FastQC version 0.12.0 (https://www.bioinformatics.babraham.ac.uk/projects/fastqc/). Reads were mapped to HfPKS using the BBmaps package version 39.01 (https://sourceforge.net/projects/bbmap/) and SAMtools version 1.18 (https://www.htslib.org/).

**Table 1.** TrapR-sequencing output and mapping statistics.

| Sequencing output | WT | PKS-IR16 | PKS-IR17 |
| --- | --- | --- | --- |
| Raw reads (demultiplexed) | 13429918 | 13528051 | 13412464 |
| Clean reads (adapter trimmed) | 13256898<br>(98,71%) | 13364326<br>(98,79%) | 13274970<br>(98,97%) |
| **Clean reads mapping to** |  |  |  |
| HfPKS ORF (6420 bp) | 1077<br>(0,01%) | 1321125<br>(9,89%) | 1382435<br>(10,41%) |
| IR stem/target site (287 bp) | 470<br>(0,004%) | 1321069<br>(9,89%) | 1381918<br>(10,41%) |
| HfPKS ORF outside IR target site (6133 bp) | 601<br>(0,005%) | 360<br>(0,003%) | 841<br>(0,006%) |
| IR spacer (117 bp) | 4<br>(0,00003%) | 2013<br>(0,015%) | 1989<br>(0,015%) |

### 2.7 dsRNA synthesis, Cy3-labeling and microscopic analysis

A 145 bp dsRNA was *in vitro* transcribed from 200 ng PCR template including opposing T7 promotors using the MEGAscript RNAi-kit (Thermo Fischer Scientific, Waltham, Massachusetts, USA). Cy3-labeling of the dsRNA was performed with the Silencer siRNA labeling kit (Thermo Fischer Scientific). A 50 µl *H. fraxineus* 1856 liquid culture was treated with approximately 140 ng/µl Cy3-dsRNA or an equivalent amount of Cy3-dye for 3 days at 22 °C. Fungal mycelium was carefully washed in 1 ml H20 without centrifugation to remove excess Cy3-dsRNA or dye, then placed onto a microscope slide and imaged with the Zeiss LSM 510 confocal laser setup connected to an Axio Observer Z1 inverted Microscope equipped with a Plan-Apochromat 20x/0.75 (Carl Zeiss, Oberkochen, Germany). Cy3 was excited at 561 nm with an argon laser and detected with a 575-615 nm bandpass filter. Image analysis was done with software ZEN 3.0 version 3.0.79.0000 (Carl Zeiss).

## 3. Results

### 3.1 Phylogenetic analysis and domain structure of RNA-interference proteins in *Hymenoscyphus fraxineus*

Despite the conservation of common RNAi components in ascomycetes, detailed identification of RNAi genes in *H. fraxineus* was unknown. Our *in silico* analysis of RNAi genes showed that homologous proteins to core enzymes of the RNAi pathway are present in *H. fraxineus* (Fig. 1). We identified six ARGONAUTE-like (AGO) proteins (Fig. 1a), two DICER-like (DCL), proteins (Fig. 1b), and additionally four RNA-dependent RNA polymerases (RDRs) (Fig 1c).

**Fig 1.**
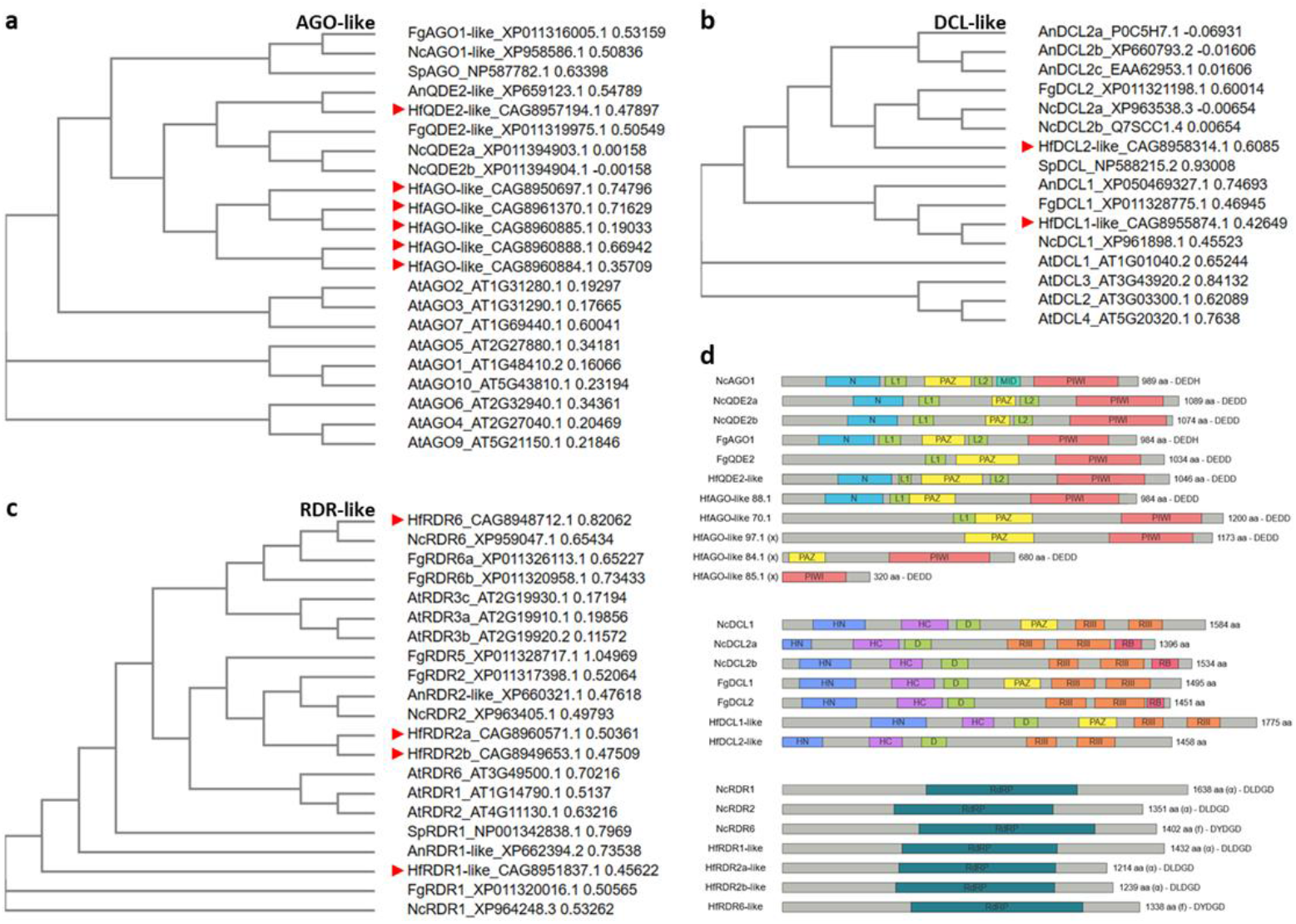
Phylogenetic trees and domain structure of RNA interference proteins. a) ARGONAUTE-like, b) DICER-like and c) RNA-dependent RNA polymerase-like genes. d) Domain structures of *Neurospora crassa, Fusarium graminearum* and putative *Hymenoscyphus fraxineus* RNAi enzymes were predicted using the InterPro database (Paysan-Lafosse et al. 2022) and visualized with the IBS2.0 webtool (Xie et al. 2022). Indicated are the catalytic tetrad DEDD/H inside the PIWI domain for AGO proteins and the signature motif DLDGD or DYDGD in the catalytic domain of RDR proteins. (x) indicates AGO-like proteins with lower domain conservation. Arrowheads indicate putative *H. fraxineus* proteins in a-c). The *A. thaliana* (At) AGO, DCL and RDR protein sequences were downloaded from TAIR (https://www.arabidopsis.org/) and used as a query in a tBLASTn search (https://blast.ncbi.nlm.nih.gov/Blast.cgi) against *Neurospora crassa* genome (Nc, NCBI:txid5141). Resulting sequences were used as query to identify homologous proteins in other fungal genomes: *Aspergillus nidulans* (An, NCBI:txid162425), *Fusarium graminearum* (Fg, NCBI:txid:229533), *Hymenoscyphus fraxineus* (Hf, NCBI:txid746836) and *Schizosaccharomyces pombe* (Sp, NCBI:txid4896). Fungal protein sequences were downloaded from the database and aligned using the EMBL-EBI webtools Clustal Omega and Simple Phylogeny (Madeira et al. 2022). N: N-terminal domain; L1, L2: linker domain; HN: Helicase domain N-terminal; HC: Helicase domain C-terminal; D: Dicer dimerization domain; RIII: Ribonuclease III domain; RB: dsRNA binding domain.

For *H. fraxineus* AGO proteins, only HfQDE-2-like retained the full set of characteristic domains present in *Neurospora crassa* AGO1 and QDE2a/b (Fig. 1d) and shared higher similarity to other fungal AGOs than to endogenous proteins. The absence of a MID domain in all except NcAGO1 suggests a lower degree of conservation in fungi. The other five HfAGO proteins clustered together, showing less similarity to fungal or plant AGO proteins. All five retained the AGO-defining PIWI domain but lost some or all other domains present in functional AGO proteins. HfAGO-like 88.1 lacks L2, and HfAGO-like 70.1 lacks both N-terminal domain and L2, similar to FgQDE2. HfAGO-like 97.1 and 84.1 only retained PAZ and PIWI domains, and the heavily truncated HfAGO-like 85.1 only consist of a PIWI domain. These three seem less likely to be functional, however we cannot rule out potential binding of an RNA and/or a target sequence. All putative HfAGO-like proteins and HfQDE2 contain the catalytic tetrad ‘DEDD’ (Asp-Glu-Asp-Asp) inside the PIWI domain. However, their function and contribution to RNAi remains unclear. Fungal DCL1 and DCL2 proteins form two separate clusters that are equally distant to *A. thaliana* DCL proteins albeit sharing critical domains like helicase and dimerization domains. Fungal DCL proteins show strong conservation of domain architecture, with a PAZ domain present in all three fungal DCL1-like proteins and a dsRNA binding domain present in Nc/FgDCL2 but not HfDCL2-like. The domain structure of RDRs in general is not well resolved, only identifying the protein family of ‘RNA-dependent RNA polymerase, eukaryotic-type’ (IPR007855). RDRs are classified by their signature motif in the catalytic domain (Wassenegger and Krczal 2006). HfRDR1 and HfRDR2a/b contain the signature motif ‘DLDGD’ (Asp-Leu-Asp-Gly-Asp) that is also conserved in fungal RDRs in this cluster and present in NcRDR1/2, and therefore belonging to α-clade. HfRDR6-like proteins, similar to NcRDR6, contain the fungal-specific RDR6 motif ‘DYDGD’ (Asp-Tyr-Asp-Gly-Asp) (f-clade) that is not present in mammals or plants (Wassenegger and Krczal 2006).

### 3.2 Stable expression of a hairpin RNA targeting a polyketide synthase-like gene

The *in silico* analysis showed the presence of putative AGO and DCL proteins in *H. fraxineus*. Both contain the characteristic domain structure found in well-studied organisms, leading to the anticipation of a functional RNAi pathway in this fungus. To investigate its functionality, *H. fraxineus* strain NW-FVA 1856 (denoted WT and/or 1856) was transformed with a construct for constitutive expression of a hairpin RNA targeting a polyketide synthase-like (PKS) gene. Fungal PKS genes are involved in melanin synthesis (Fig. 2a) and accumulation of melanin leads to a dark colouration of mycelium. (Takano et al. 1995; Akamatsu 2010). Two independent transgenic lines were regenerated which displayed the expected albino-like phenotype (Fig. 2b). The lack of dark pigmentation in the PKS-IR expressing lines indicates functional silencing of the PKS gene. Analysis of PKS-transcript abundance relative to UBC showed no reduction of mRNA levels in both IR-expressing lines when compared to the wild-type (Fig. 2c). The cDNA template was synthesized from total RNA extracts using oligo(dT) primers. Subsequently, PKS-specific primers amplified fragments of the cDNA that were upstream (5’) or downstream (3’) of the PKS-IR target site in the PKS ORF (Fig. 2d). Presence of both amplicons indicated that the cDNA template and presumably also the mRNA was intact over the analysed region.

**Fig 2.**
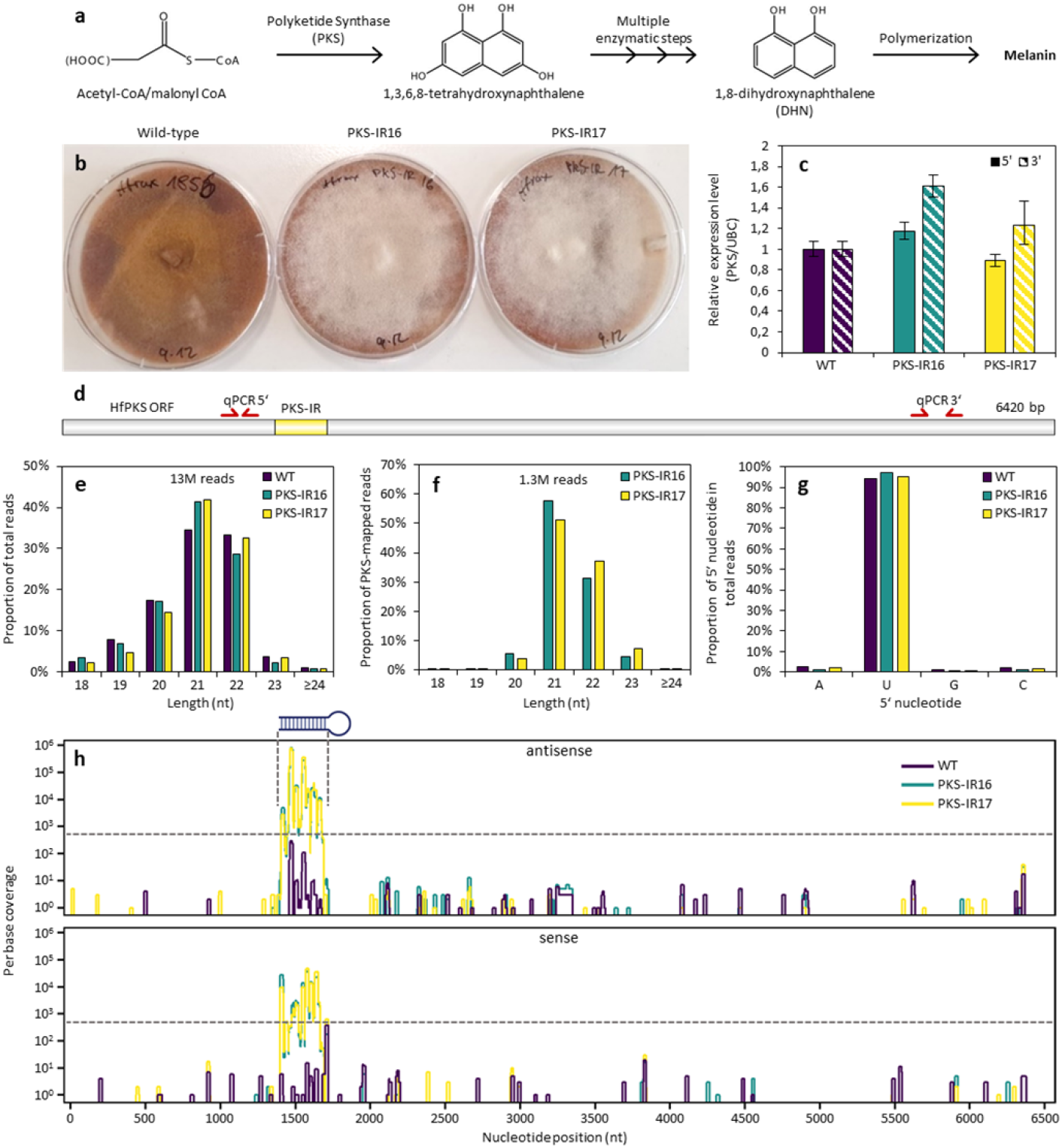
Stable expression of a hairpin RNA targeting a polyketide synthase-like gene in *H. fraxineus*. a) Acetly-CoA/malonyl-CoA is the substrate for PKS in the biosynthesis pathway of melanin (adapted from Eisenman and Casadevall 2012). b) Albino-like phenotype in PKS-IR expressing transgenic lines. c) Relative expression level of PKS relative to UBC reference. d) Location of qPCR primer pairs and PKS-IR target site in HfPKS ORF. Processing of PKS-IR was analysed by small RNA sequencing of AGO-bound small RNAs using the TrapR-kit (Lexogen). e) Length distribution of total reads. f) Length distribution of reads mapping to HfPKS. g) 5’ nucleotide content in total reads. h) PKS-IR derived reads mapping to the stem of PKS-IR. Dashed line indicates background contamination threshold at 500 reads.

The processing of the PKS-IR RNA was analysed by extracting the RNA-induced silencing complex (RISC) from fungal mycelium using Lexogen’s TrapR-Kit and Illumina sequencing. The method is based on conserved structural and biochemical properties of AGO proteins, that utilise ion exchange chromatography to separate contaminating nucleic acids (e.g. degradation products) from RISC-containing cell-lysate extracts (Grentzinger et al. 2020). Extraction of AGO-bound siRNAs from the RISC enriched fraction enables the analyses of biologically relevant small RNAs.

Subsequently, siRNAs were subjected to small RNA sequencing that detected approximately 13 million total reads in each sample for WT and both PKS-IR expressing lines (Fig. 2e). In *H. fraxineus*, small RNAs with a length of 21- and 22-nt were enriched in WT, PKS-IR16 and PKS-IR17 samples, accounting for 68%, 70% and 75% of total reads, respectively. However, 21- and 22-nt siRNAs were equally abundant (35% and 33% of total reads, respectively) in the wild-type, 21-nt siRNA abundance was increased by 13% and 9% in PKS-IR16 and PKS-IR17, respectively, when compared to 22-nt siRNA abundance. The enrichment of 21- and 22-nt siRNA was even more pronounced when looking at PKS-IR16 and PKS-IR17 reads mapped to the HfPKS open reading frame (Fig. 2f). PKS-mapped reads made up 10% of total reads in both PKS-IR samples. Interestingly, all samples showed a strong bias for uracil (U) as the 5’ terminal nucleotide with more than 90% compared to other bases in all detected reads (Fig. 2g). In both transgenic PKS-IR expressing lines reads showed sequence homology in high abundance (1.3 M reads of 13 M total) to the double-stranded stem region of the PKS-IR sequence, with a precise boundary at the first and last nucleotide of the 287 bp stem (Fig. 2h). Reads mapping to the antisense strand were enriched over reads in sense orientation, possibly due to stabilization of guide strands, which are protected by AGO homologues in *H. fraxineus*. A negligible number of reads mapped to the HfPKS ORF outside of the IR region, and in the wild-type sample only 1077 reads mapped to HfPKS ORF.

This analysis suggests that the processing of the PKS-IR RNA in *H. fraxineus* follows the general model of RNAi, first cleaving a long dsRNA into siRNA and loading it into AGO. However, qRT-PCR results suggest that the siRNAs do not lead to cleavage of the target RNA as in plants.

### 3.3 Exogenous application of fluorescently labelled dsRNA to *H. fraxineus* liquid culture

Since exogenous application of dsRNA has already shown promise as a plant protection strategy, we wondered whether *H. fraxineus* would readily take up fluorescently labelled dsRNA. Using confocal laser scanning microscopy of untreated fungal mycelium, no background fluorescence was detected in the Cy3-channel (Fig. 3a-c), whereas in samples treated with the free Cy3 dye, a fluorescent signal was detected in some hyphae, localised to small round structures (Fig. 3d-i). After incubating fungal mycelium in liquid culture for 3 days with approximately 140 ng/µl of Cy3-dsRNA, fluorescent signals following the internal structures of the hyphae were observed (Fig. 3j-o). The Cy3 signal colocalised with densely packed structures, as well as a more diffuse signal in less densely packed areas, and in some cases omitted clear spaces that appear to be vacuoles. Since other fungal cell compartments are situated in the cytoplasm, it seems plausible that the Cy3 dsRNA also entered the cytoplasm.

**Fig 3.**
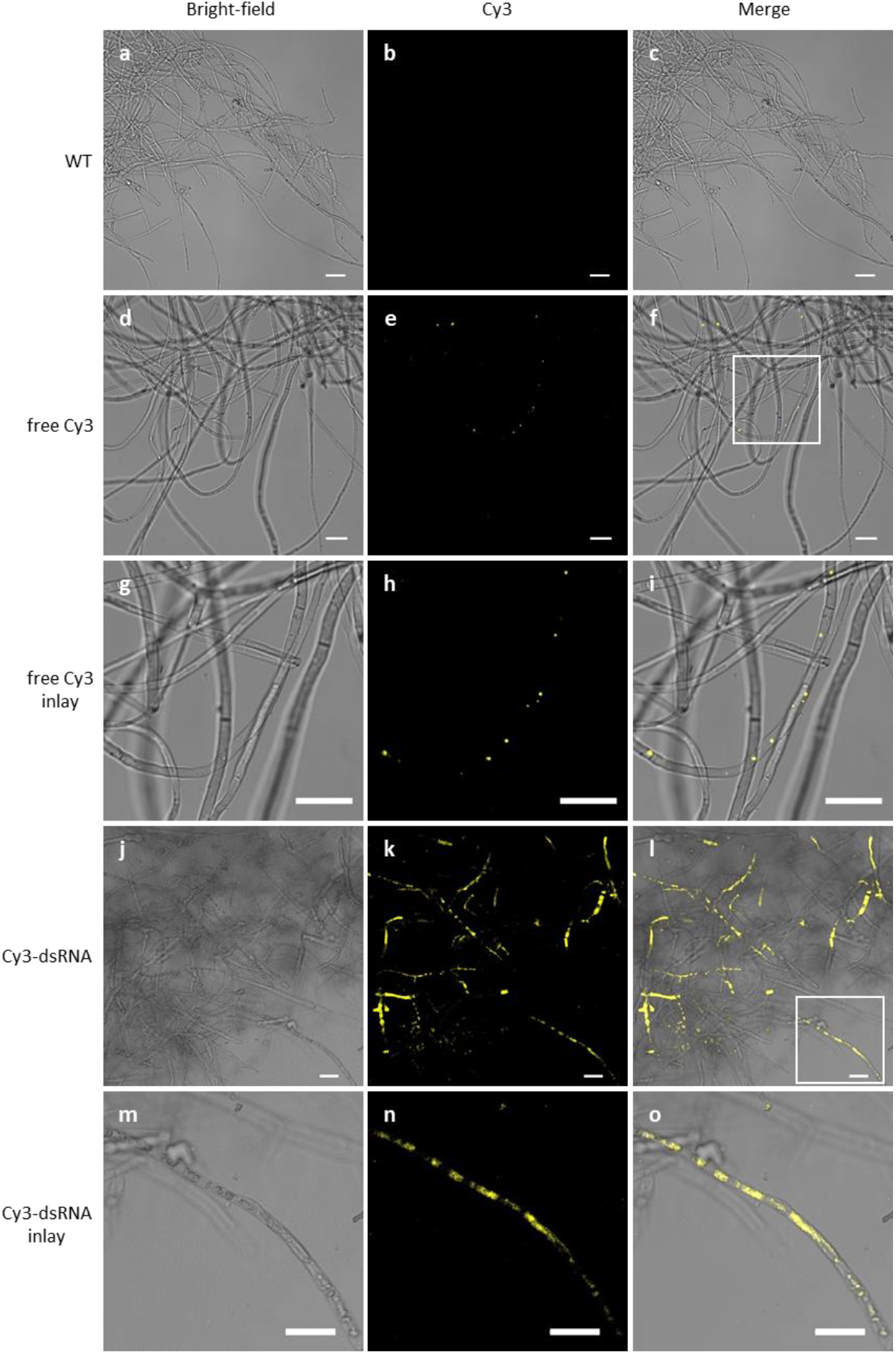
Uptake of exogenously delivered fluorescently labelled dsRNA by *H. fraxineus*. a-c) wild-type mycelium of an *H. fraxineus* liquid culture showing no background fluorescence. d-f) Cy3-dye treated samples showed small dots of fluorescence that colocalized with dark round structures of the fungal hyphae. g-i) Inlay of f). j-l) Cy3 fluorescent signals following internal structures of fungal hyphae were detected after 3 days of incubation with approximately 140 ng/µl Cy3-labelled dsRNA. All scale bars = 20 µm

## 4. Discussion

In this study, we provided the first evidence for a functional RNA interference mechanism in *H. fraxineus* by analysing the albino-like phenotype observed in transgenic fungal lines expressing a hairpin RNA targeting a polyketide synthase-like gene.

Results from qRT-PCR analysis indicated that gene silencing was caused by translational inhibition rather than transcript cleavage or DNA methylation. The cDNA template was synthesised from total RNA extracts using oligo d(T) primers. PKS-specific primer pairs were designed to bind upstream (5’) and downstream (3’) of the PKS-IR target site. Both pairs generated amplicons in two PKS-IR lines that were not reduced but rather equal or slightly more abundant compared to the wild-type sample and relative to the reference gene. Cleavage would occur at the IR-target site, thus it would not have been possible to synthesize cDNA and amplify the 5’ upstream region. The qRT-PCR analysis also indicated that gene silencing at the transcriptional level seems less likely as a cause for the phenotype, since DNA methylation or heterochromatin would have prevented the initial synthesis of the mRNA leading to reduced transcript levels (Erdmann and Picard 2020).

Sequencing of active AGO-bound, PKS-IR derived primary siRNAs were mapped exclusively to the stem of the IR and no secondary siRNAs were detected. The formation of secondary siRNA was not widely reported in other fungal species (Mann et al. 2023). However, Kowalik et al. (2015), who analysed the expression of a hairpin RNA in the fission yeast *Schizosaccharomyces pombe* showed that the formation of secondary siRNAs was negatively regulated by the RNA polymerase-associated factor 1 complex (Paf1C). Similar to our results, a stably expressed hairpin RNA was cleaved into primary siRNAs without reducing transcript levels of the targeted gene. When the activity of Paf1C was impaired by introducing premature stop mutations in subunits of the complex, secondary siRNAs were detected in high abundance covering the whole transcript, leading to transcript downregulation and *de novo* formation of stable heterochromatin. *In silico* analysis identified homologous proteins for five out of six subunits of Paf1C in *H. fraxineus* (Paf1, CAG8954017.1; Cdc73, CAG8957552.1; Prf1, CAG8949795.1 and Tpr1, CAG8958833.1 but not Leo1). Future experiments should be designed to investigate the function of the Paf1 complex and decipher the potential interplay with RNAi enzymes in *H. fraxineus*.

MicroRNA (miRNA)-mediated post-transcriptional gene regulation has been extensively studied in plants and animals. MiRNA hairpin precursors are transcribed from endogenous loci with subsequent processing in the canonical RNAi pathway, yielding mature miRNA that contain mismatches with the target sequence. The degree of homology to the target transcript determines the regulatory mechanism (either cleavage or blockage) by RISC (Moran et al. 2017). Both mechanisms are present in both kingdoms, but target cleavage is more common in plants and translational repression is more common in animals. Plant miRNAs require high sequence homology to cleave the target transcript efficiently, conferring high specificity to a small number of transcripts. In animals, miRNA target recognition is initiated by the 7 nt seed region (position 2-8 in mature miRNA), leading to translational repression through inhibition of translation initiation, elongation or deadenylation, thus potentially regulating a large number of genes (Moran et al. 2017). In both pathways, a GW-rich protein (GW148/TNRC6 in animals and SUO in plants) is recruited to RISC to mediate translational repression. The class of GW-rich proteins shows very limited sequence homology and class-defining domains have yet to be determined, therefore no homologous proteins were identified in *H. fraxineus* and other fungi (species analysed as in Fig. 1). The regulatory differences highlight the need for in-depth analysis of the mechanisms involved in translational repression in fungi.

AGO-bound siRNAs implicate special features in *H. fraxineus*. In *Neurospora crassa*, recombinantly expressed dsRNAs are cleaved primarily into 25-nt siRNAs (Li et al 2010), whereas the most prominent cleavage products of PKS-IR in *H. fraxineus* are 21- and 22-nt long. In *N. crassa*, 20-21nt siRNAs originate from DNA damage-induced dsRNAs (Lee et al. 2009). Despite differences in size, 5’ nucleotide enrichment in *H. fraxineus* is very similar to other fungi. AGO-bound siRNAs in *H. fraxineus* have 5’ uracil enrichment irrespective of its origin, PKS-IR or endogenous sequences. In *N. crassa*, miRNA-like sRNAs, which require canonical RNAi components, or 22-nt long siRNAs, which originate from overlapping transcripts and do not require canonical RNAi proteins, show 5’uracil enrichment (Lee et al. 2010). Similarly, exonic-siRNAs in *Mucor circinelloides* are predominantly 21-nt or 25-nt long and contain a 5’uracil (Cervantes et al. 2013). In plants, the 5’ nucleotide determines the sorting of siRNAs into different AGO proteins, facilitating their diverse mechanisms in gene regulation (Mi et al. 2008). This characteristic is generated by the cleavage position of DCL proteins and more specifically, AtDCL4 preferentially generates 5’ uracils (Vaucheret and Voinnet 2023), whereas siRNAs produced by AtDCL3 are enriched with 5’ adenines (Wang et al. 2023). Shared 5’ uracil prevalence between *H. fraxineus* and other fungi but the discrepancy in the size may also be due to the differences in the sRNA-sequencing methods. Using TrapR, we cannot rule out that 25-nt siRNAs are not captured. However, our results clearly demonstrate that the amount of AGO-bound 25-nt siRNAs is negligible in *H. fraxineus*.

The decolouration phenotype in *H. fraxineus* containing the PKS-IR and the siRNA signature suggests the presence of functional RNAi machinery in *H. fraxineus*. A promising approach for controlling fungal infections is using exoRNAs (Dalakouras et al. 2020, Degnan et al. 2023, Koch et al. 2016). An important step for the efficacy of exogenous RNA delivery is the uptake of dsRNAs into the fungal cells. Using confocal microscopy, we determined that dsRNAs were taken up by the hyphae. However, there are two major aspects for further improvement. The first one is the ratio of hyphae, which takes up dsRNA as we detected Cy3-tagged dsRNA only in a limited number of hyphae. The second one is the distribution of the Cy3-tagged dsRNA signal indicating two distinct localizations of the dsRNA. The spotty localization, which is partly shared in Cy3-only control and the wide cytoplasmic signal (Fig. 3n, 3o). Unlike the cytoplasmic signal in rust fungi (Degnan et al. 2023), the Cy3-dsRNA does not cover the whole hyphae, either due to structural differences or limited dsRNA uptake efficiency in *H. fraxineus*.

Another crucial step for using the dsRNA to fight ash dieback is determining a target gene. While PKS give a visible, measurable readout for RNAi efficiency, the aim is to target gene(s), which are vital for *H. fraxineus*. Previous studies demonstrated that a number of different genes could serve as targets: 1) The pustules formed on *Syzygium jambos* whole plants by the rust fungus *Austropuccinia psidii* were strongly reduced when plants were inoculated with a mixture of urediniospores and 100 ng/µl dsRNA targeting fungal 28S ribosomal RNA, translation elongation factor 1-α or β-tubulin. At two weeks post-inoculation, the diseased leaf area was reduced by 81%, 95% and 100%, respectively, compared to no-target dsRNA control (Degnan et al. 2023). 2) Detached barley leaves were sprayed with 20 ng/µl dsRNA targeting three cytochrome P450 sterol 14α-demethylase (CYP51A, B, C) genes separately in *F. graminearum* that strongly reduced infected leaf area by 78-82% and increased to 93% reduction when all three genes were targeted at once in a single dsRNA (Koch et al. 2019). 3) In a screening of 59 target genes in *Sclerotinia sclerotiorum*, 20 genes were found to reduce the lesion size on *Brassica napus* leaves when targeted with topically applied dsRNA (McLoughlin et al. 2018). The highest reduction in lesion size detected was 85% when targeting SS1G_01703 or SS1G_06487, however on *A. thaliana* leaves the reduction was 46% and 66%, respectively.

Targeting the homologue of SS1G_06487 in *Botrytis cinerea* (BC1G_04775), necrosis on *B. napus* leaves was reduced by 53%. The strongest reduction (66%) of *B. cinerea* lesion size on *B. napus* leaves was achieved by targeting BC1G_04955. On the other hand, the *S. sclerotiorum* homologue SS1G_02495 showed strong alterations in efficiency (*A. thaliana*: 36%, *B. napus*: 71%). 200 ng or 500 ng dsRNA were topically applied to each leaf in infection assays for *S. sclerotiorum* and *B. cinerea*, respectively. These studies demonstrate the great potential of RNAi-based plant protection strategies. However, a critical evaluation of selected target genes in infection assays is essential and target prediction should not solely rely on *in silico* analysis of homologous genes. Multiplexing target genes in various combinations to a single dsRNA molecule is a potential solution to increase silencing efficiency, even for single genes with already good efficiency.

After spraying, dsRNA does not persist long on the surface of plants. There are chemical additives, which are shown to increase the half-life of dsRNA. Carbon nanoparticles synthesised from glucose and polyethyleneimine were used to deliver siRNAs into *Nicotiana benthamiana* and *Solanum lycopersicum* plant cells, effectively silencing a transgene (GFP) and endogene (Magnesium Chelatase, only *N. benthamina*) (Schwartz et al. 2020). The nanoparticle-associated RNA was protected against RNase degradation for at least 60 min. However, the ability of this carrier to confer plant protection against fungal diseases remains to be elucidated. One promising carrier molecule is layered double hydroxide, a chemical structure found naturally in minerals, which, when complexed with dsRNA targeting two (DCL1 and DCL2) or three (VPS51, bik1 and SAC1) genes in *B. cinerea*, has been shown to increase the duration of protection against *B. cinerea* infection from 5 to 10 days on tomato fruits and from 1 to 3 weeks on tomato leaves (Niño-Sánchez et al. 2022).

Since *H. fraxineus* grows in the xylem, stem injection and petiole feeding may be promising approaches to control ash dieback. Using these methods for the application of Cy3-labelled dsRNA, a fluorescent signal was detected in the xylem and apoplast of leaves of *Malus domestica, Vitis vinivera* and *Nicotiana benthamina* (Dalakouras et al. 2018). The petiole-feeding method may be more appropriate for treating smaller plants, as reaching the canopy of large trees can be difficult. On the other hand, trunk injection can be scaled up to the size of the tree with repeated applications or by using a large container reservoir. To ensure resource efficiency, it is important to time any RNA-based application method in relation to the deposition of airborne ascospores by *H. fraxineus*. Trunk injection is a suitable method for responding quickly to high spore loads. Applied RNA can be detected in leaves’ veins as soon as one day after application, as demonstrated by Dalakouras et al. (2018).

## Conclusion

Our work provides the essential basis, specifically the presence of RNAi pathway in *H. fraxineus* and the uptake of exogenously applied dsRNA, for the development of an RNAi-based approach to protect ash trees from the devastating disease ash dieback caused by this fungus. Existing methods for exogenous delivery of dsRNA are well suited to targeting the fungus at different stages of its life cycle. The primary aim of future research should be to investigate the molecular mechanisms that underlie RNA interference and the potential suppressors of silencing in *H. fraxineus*.

## Acknowledgement

The authors thank Moritz Hohenwarter for artistic suggestions to the figures and Max Preusse for improving the manuscript with edits for language, grammar, and clarity.

## Supplementary Information

Supplementary Table S1 List of primers used in this study.

Supplementary Fig. S1 Vector map of pSilent PKS-IR

