## Supplementary Table S1 and Fig. S1 for "Functional RNA Interference in *Hymenoscyphus fraxineus:* Hairpin RNA-Induced Gene Silencing Of A Polyketide Synthase-like Gene": Hohenwarter_etal_supplementary information.docx

**Journal of Plant Diseases and Protection**

Linus Hohenwarter^1^, Arvid Hanke^1,4^, Birgit Hadeler^2^, Alexandra Bassler^1^,
Gitta Langer^3^, Gabi Krczal^1^, Veli Vural Uslu^1,4,*^

**Affiliations**

^1^Translational Epigenetics, RLP Agroscience GmbH, Breitenweg 71, D-67435 Neustadt-Mussbach, Germany

^2^Institute of Plant Science and Microbiology, Molecular Phytopathology, University of Hamburg, Ohnhorststrasse 18, D‑22609 Hamburg, Germany

^3^Department of Forest Protection, Northwest German Forest Research Institute (NW‐FVA), Grätzelstraße 2, D‐37079 Göttingen, Germany

^4^Centre for Organismal Studies (COS) Heidelberg, University of Heidelberg, INF230, D‑69117 Heidelberg, Germany

**Table S1**: Primer used in this study

| **Primer ID** | **Sequence** | **Purpose** |
| --- | --- | --- |
| pSilent1_insert_fw | CAAGCATCGATACCGTCG | Genotyping for presence of PKS-IR, separate amplification of sense and antisense arm |
| Spacer_rv | GTAACGTTAAGTGGATCCGG |  |
| Spacer_fw | GTCAAATTGATTCTCGGGAAGC |  |
| pSilent1_insert_rv | GCTTCCCGAGAATCAATTTGAC |  |
| Hygro-fw | CGGCCGCGCTCCCGATTC | Genotyping for presence of hygromycin resistance cassette |
| Hygro-rv | CCAGAAGAAGATGTTGGCGACC |  |
| HfPKS_fw1 | CTTTCAAGTCAACGCCAAC | qPCR of polyketide synthase like gene, 5’ fragment |
| HfPKS_rv1 | GTATGCAAGGGCAGCAG |  |
| HfPKS_fw2 | CTTATCTTCTCGCAGGCTG | qPCR of polyketide synthase like gene, 3’ fragment |
| HfPKS_rv2 | CTTCATCATGCTGAAGTGG |  |
| HfUBC_ fw | CAGACACCCCATTTGAAG | qPCR of reference gene UBC2 |
| HfUBC_ rv | GTGGCATATACGTTTGGATG |  |

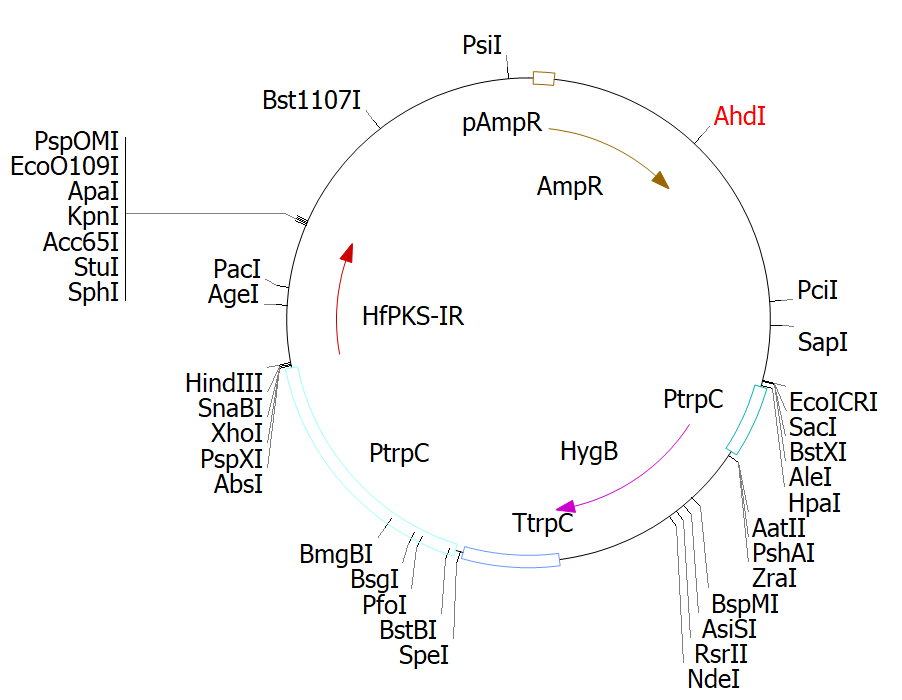

**Fig. S2** Vector map of pSilent PKS-IR
